# Does media matter? Growth environment influence antimicrobial tolerance and expression of virulence and transmembrane ion transport-associated genes in MRSA

**DOI:** 10.64898/2026.06.15.732343

**Authors:** Oluwatosin Qawiyy Orababa, Ayomikun Kade, Leonard Ighodalo Uzairue

**Affiliations:** School of Life Sciences, Gibbet Hill Campus, University of Warwick, Coventry, CV47AL, United Kingdom; School of Allied Health Sciences, Faculty of Health and Life Sciences, De Montfort University, The Gateway, Leicester, LE1 9BH, United Kingdom

**Keywords:** MRSA, growth environment, AMR, transmembrane transport

## Abstract

Clinically relevant pathogens are often tested for antimicrobial susceptibility using standard laboratory media that poorly reflect the *in vivo* environments in which they cause infections, leading to poor clinical outcomes. In this study, we aim to understand the impact of media on the global transcriptome, biofilm formation, and antibiotic susceptibility of methicillin-resistant *Staphylococcus aureus* USA300 when cultivated in a physiologically relevant wound medium, such as simulated wound fluid (SWF), compared to cation-adjusted Mueller-Hinton broth (caMHB), a general-purpose medium. The transcriptomics analysis showed upregulation of 865 genes and downregulation of 792 in SWF compared to caMHB. Upregulated genes in SWF are associated with virulence, such as genes coding for fibronectin-binding proteins (*fnaAB*), serine proteases (*splABCDE*), as well as genes involved in antimicrobial resistance, such as multidrug efflux pump genes *(norB, norC*). Conversely, genes associated with transmembrane ion transport, including phosphate transport (*pstSCAB, phoU*) and potassium intake (*kdpABCF*), were significantly downregulated in SWF, as further confirmed by increased membrane disruption upon exposure to a membrane-potential-sensitive dye (DiSC3). Biofilm assay showed reduced surface attached biofilm but increased cell-to-cell attachement in SWF compared to caMHB. Antimicrobial susceptibility testing revealed a 2- to 4-fold increase in tolerance to clinically relevant antibiotics in SWF compared to caMHB. Overall, our findings revealed that media affects gene expression, membrane physiology, virulence, and antibiotic tolerance in MRSA, underscoring the need to use physiologically relevant media in routine antimicrobial susceptibility testing and the drug development pipelines.

## Introduction

Around 5 million deaths are currently associated with antimicrobial-resistant (AMR) infections, globally (Naghavi et al., 2024). Oftentimes, some of these deaths involve pathogens that have *in vitro* susceptibility to current antimicrobials; however, they become highly tolerant to these antimicrobials *in vivo*, leading to worse clinical outcomes (Taubenberger et al., 2026). This variation between *in vitro* and *in vivo* susceptibilities is often attributed to environmental factors that increase antimicrobial tolerance in bacterial pathogens in their natural growth environments (Anbo et al., 2026). Many growth media used for either preclinical antimicrobial testing or clinical antimicrobial susceptibility testing do not provide conditions similar to those found *in vivo*, which accounts for the inconsistencies observed between *in vitro* and *in vivo* activities. Hence, to effectively study and understand bacterial pathogenicity and antimicrobial susceptibility, it is important to provide conditions that mimic *in vivo* infection environments. This is especially important during preclinical testing of new antimicrobials and during clinical diagnosis.

In this study, we mainly used transcriptomics to understand the variations in gene expression of *Staphylococcus aureus* USA300, a methicillin-resistant (MRSA) strain, in two growth environments: cation-adjusted Mueller-Hinton broth (caMHB) and simulated wound fluid (SWF), representing standard laboratory media and growth-mimicking media, respectively. The choice of MRSA in this study is due to its status as a high priority pathogen for which new drugs are required (Sati et al., 2025). This pathogen also accounts for about 1 million annual AMR-associated deaths globally, which is highest among bacterial pathogens (Naghavi et al., 2024). Additionally, in 2019, *S. aureus* infections were linked to a productivity loss of approximately $36 billion, most of which was due to MRSA infections (Naylor et al., 2025). MRSA is also implicated in chronic infections, including chronic wound infections and cystic fibrosis lung infections, due to its ability to form robust biofilms in these infection environments (Kaushik et al., 2024; Touaitia et al., 2025). Hence, to effectively study this pathogen and develop treatments for its infections, it is important to use the right growth environment/model.

This study revealed strong variation in the gene expression of *S. aureus* USA300 across the two growth environments (caMHB and SWF), including decreased expression of transmembrane ion transport-associated, efflux, and virulence-associated genes. This emphasises the need to provide the right growth environment during preclinical analysis as well as clinical diagnosis.

## Result

### Variations in the global transcriptome of MRSA in caMHB and SWF

To understand the level of variation in the global transcriptome of *S. aureus* USA300 (a methicillin-resistant strain) in different growth environments, we performed a differential gene expression (RNA sequencing) experiment, comparing cultures of *S. aureus* USA300 in cation-adjusted Mueller-Hinton Broth (caMHB) and synthetic wound fluid (SWF) at 6h of growth. A total of 865 and 792 genes were significantly upregulated and downregulated in SWF compared to caMHB, respectively (Figure 1a), indicating substantial variation in the significantly expressed genes in S. aureus across both media. Both volcano plots and heatmaps of significantly expressed genes in SWF compared to caMHB further reveal strong variations in expression pattern (Figure 1b and 1c).

**Figure 1.**
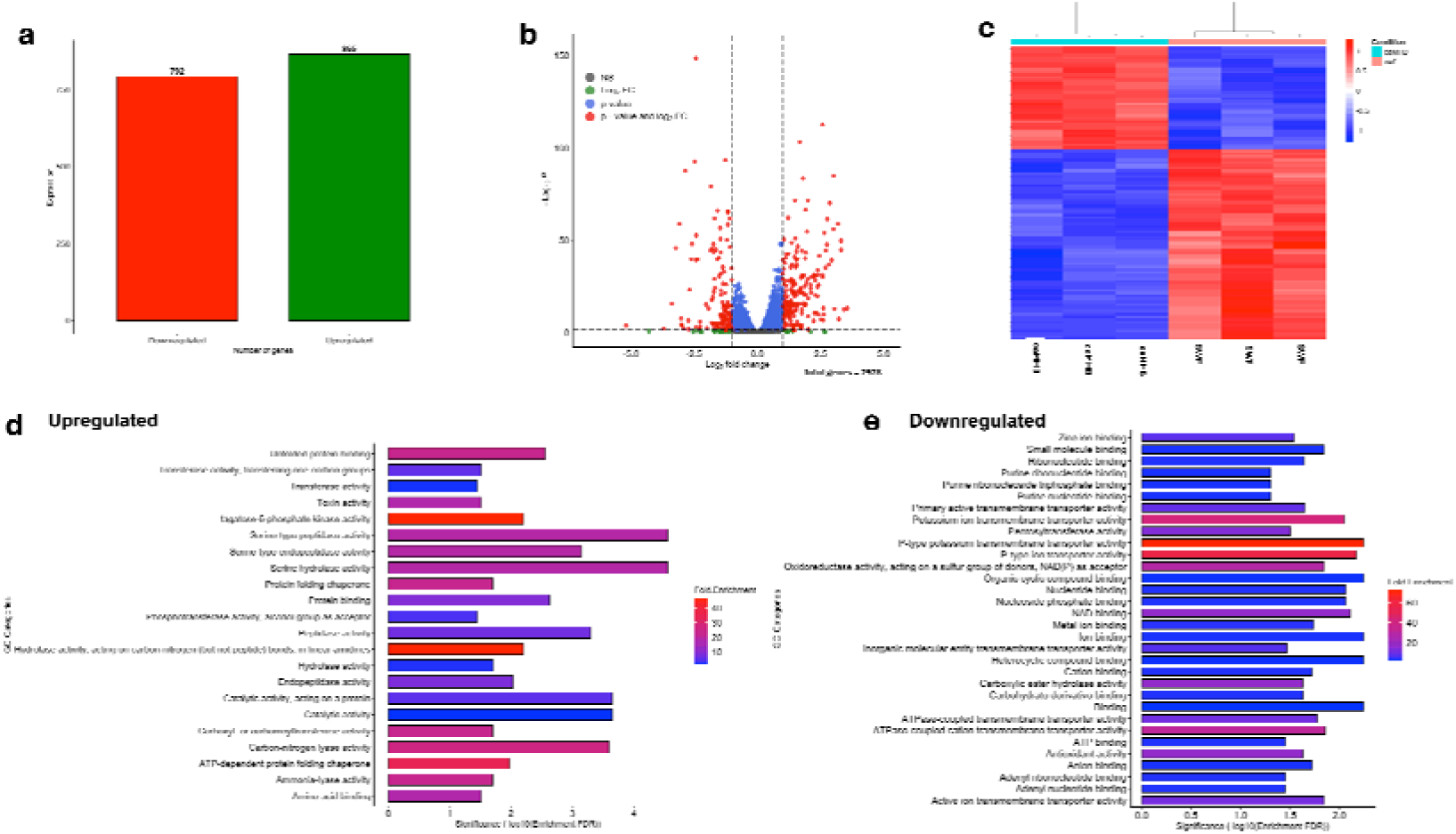
Significant variation in MRSA global transcriptome in caMHB and simulated wound fluid. a. Number of significantly (p<0.05) upregulated and downregulated genes in SWF, when compared to caMHB. b. Volcano plot showing differentially expressed genes (p<0.05 and |log2foldchange| ≥ 1) in red. Heatmap of the top 100 differentially expressed genes in SWF and their pattern of expression in both media. d. Differentially upregulated gene ontology groups (molecular function) in SWF compared to caMHB e. Differentially downregulated Gene Ontology (Molecular Function) groups in SWF compared to caMHB.

Of great interest among the significantly upregulated gene ontology groups in SWF compared with caMHB are those linked to toxin production, serine-type peptidase, serine-type endopeptidase, and serine hydrolase activities (Figure 1d). On the other hand, majority of the downregulated GO groups were associated with transmembrane ion transport. This includes GO groups associated with anion binding, cation binding, potassium ion transmembrane transporter, metal binding, inorganic molecular entity transmembrane transporter, and zinc ion binding (Figure 1e). Among the significantly downregulated genes, the *kdpABCF* genes are primarily associated with potassium uptake across the plasma membrane (Figure 2a-b). Genes involved in phosphate transport (*phoU* and *pstABCS*) were also differentially upregulated in SWF compared to caMHB (Figure 2a). These genes work together to transport phosphate into the cell, as shown in Figure 2b. The major genes highlighted under the serine-type endopeptidase/peptidase groups are the serine proteases (*splABCDE*) (Figure 2b). These serine protease-like genes have been previously associated with *S. aureus* pathogenesis and immune invasion (Pospich et al., 2026; Dasari et al., 2022). We also observed the upregulation of *fnbA* and *fnbB* (fibronectin-binding proteins), *hlgBC* (bicomponent toxins), and alpha-phenol-soluble modulins (Figure 2c). Another interesting expression pattern observed is the significant upregulation of multidrug efflux pump genes, including *norB* and *norC*, which belong to the major facilitator superfamily (Figure 2a-b).

**Fig 2.**
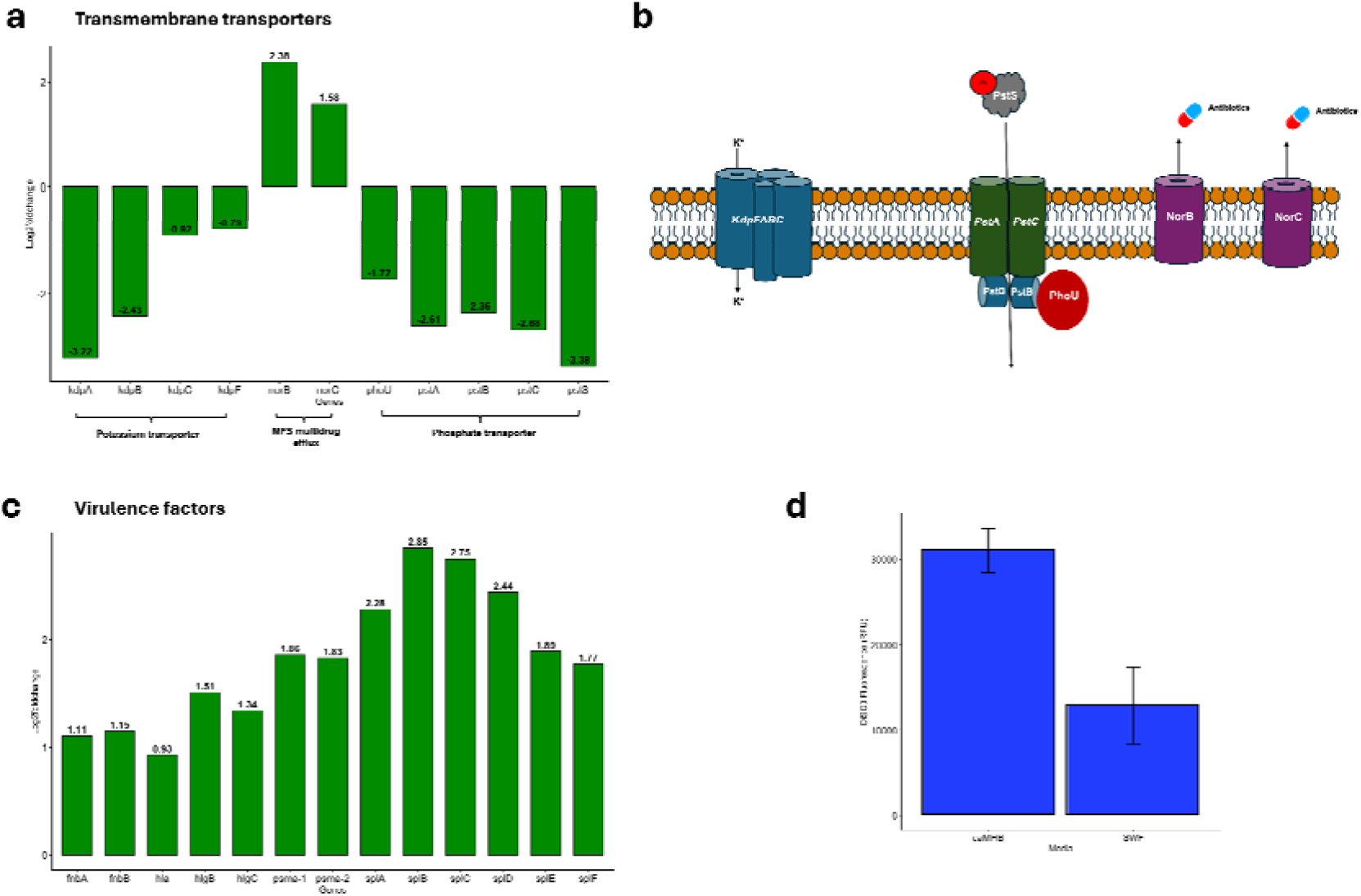
Decreased transmembrane transport activity and increased expression of virulence genes in simulated wound fluid. **a.** Expression patterns of differentially expressed transmembrane transport proteins in SWF compared to cation-adjusted Mueller-Hinton broth (caMHB). **b.** A membrane model showing the function of some differentially expressed transmembrane proteins. **c.** Differential upregulation of virulence factors in MRSA in SWF. **d.** DiSC3(5) fluorescence showing membrane activities in both caMHB and SWF. S. aureus USA300 was stained with DiSC3(5) and end fluorescence measured after 1 h. Error bars represent biological triplicates (N= 3).

### Decreased expression of membrane transport genes and increased expression of virulence genes in SWF compared to caMHB

Due to the significant upregulation of gene ontology groups associated with transmembrane transport, including potassium and phosphate transports (Fig 1), and the role of potassium in maintaining stable bacterial membrane potential. We further zoomed into the genes associated with the activities and their expression levels. We observed the upregulation of all the genes in the kdpFABC (potassium transport activity) and the *pstSCAB*-*phoU* operons (phosphate transport) (Fig 2a-b). Genes coding for multidrug efflux transporters (*norB* and *norC*) in *S. aureus* were also significantly upregulated (Fig 2a-b). Additionally, due to the upregulation of the GOs associated with toxins and serine proteases, we zoomed into these genes and looked at those that were significantly upregulated. All genes in the *spl* operon (serine protease-like protein coding operon) were significantly upregulated as well as notable *S. aureus* toxin coding genes including alpha hemolysin (*hla*), gamma hemolysin (*hlg*), and phenol-soluble modulin alpha (*psm-a*). Considering the importance of potassium and phosphate transport in maintaining a stable bacterial membrane potential, we decided to explore how these two growth environments influence bacterial membrane potential. To do this, we used the membrane-potential-sensitive dye DiSC3, whose fluorescence increases with greater disruption of the membrane potential. *S. aureus* USA300 LAC was exposed to DiSC3, and end fluorescence was measured after 30 min. We observed a significant increase in DiSC3 fluorescence in MRSA in caMHB compared to SWF, further supporting the transcriptomics evidence of decreased expression of genes associated with ion transmembrane transport in SWF.

### MRSA has different biofilm morphology wound-mimicking environment

To understand how different growth environments might influence biofilm formation, we grew *S. aureus* USA300 in a caMHB and SWF in a 48-well plate for 48h. After 24h of growth, we observed the growth morphology of this bacterial strain in both media and noted some form of cell clumping in SWF but not in caMHB (Fig 3a). After 48 h, we measured biofilm formation using the crystal violet staining technique and, interestingly, observed lower biofilm yield in SWF compared to caMHB (Figure 3b-c). This might possibly be due to the loosely attached biofilm in SWF that were washed away. Lastly, we carried out fluorescent imaging with calcofluor white in a soft-tissue collagen wound model that has been previously reported to mimic an actual wound environment. We observed that MRSA forms micro-colonies in this soft tissue wound model (Figure 3d), which possibly explains the morphology (clumps) observed in SWF in Figure 3a. This microcolony morphology in wound infections has also been previously reported (Davis et al., 2008).

**Figure 3.**
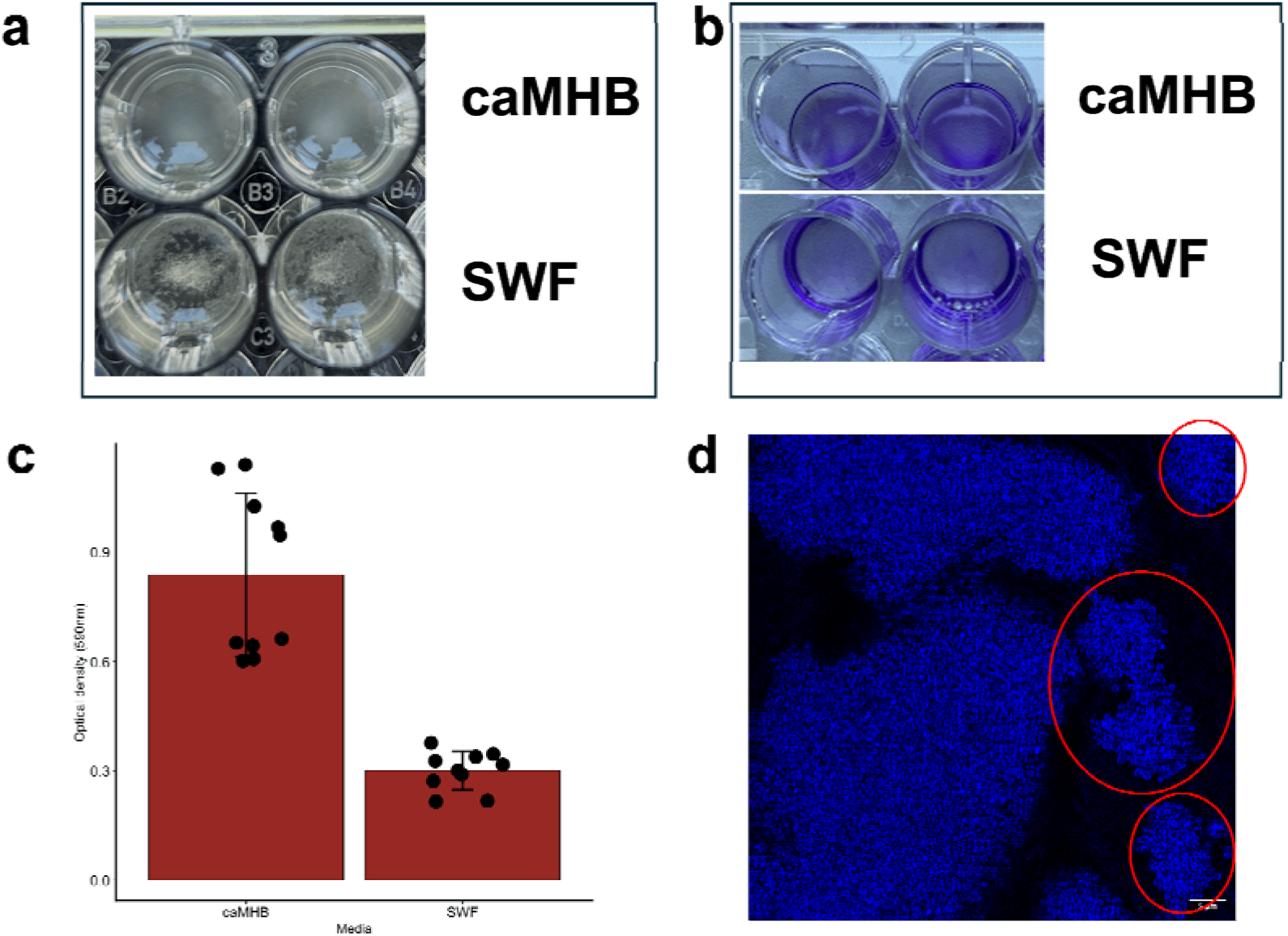
Morphological changes in MRSA biofilm across both media. **a.** Growth morphology of *S. aureus* USA300 LAC in caMHB and SWF after 24 h of growth in a 48-well plate. **b.** Crystal violet staining of 48-h biofilm of *S. aureus* USA300 LAC in both caMHB and SWF. **c.** Biofilm biomass (at OD590nm) of 48h biofilm of c in both caMHB and SWF after crystal violet staining. **d.** Confocal fluorescent microscopy showing the formation of microcolonies by *S. aureus* USA300 in a soft tissue wound biofilm model.

### MRSA shows higher tolerance to clinically relevant antibiotics in SWF compared to caMHB

Due to increased expression of efflux genes in SWF and the reduced transmembrane transport activity (likely causing reduced antibiotic uptake), we hypothesised that MRSA would be more resistant to antibiotics in SWF compared to caMHB. To test this hypothesis, we performed antimicrobial susceptibility testing of some clinically relevant antibiotics against S. aureus USA300 LAC using the broth microdilution assay. Overall, we observed that all tested antibiotics had higher minimum inhibitory concentrations (MICs) in SWF than in caMHB (Figure 4). Interestingly, this variation was more pronounced at the bactericidal level (minimum bactericidal concentrations – MBC) (Figure 4). We observed twofold higher MBC values in SWF than in caMHB. This increase in MIC and MBC values might be associated with increased efflux gene expression or decreased transmembrane transport activity.

**Figure 4.**
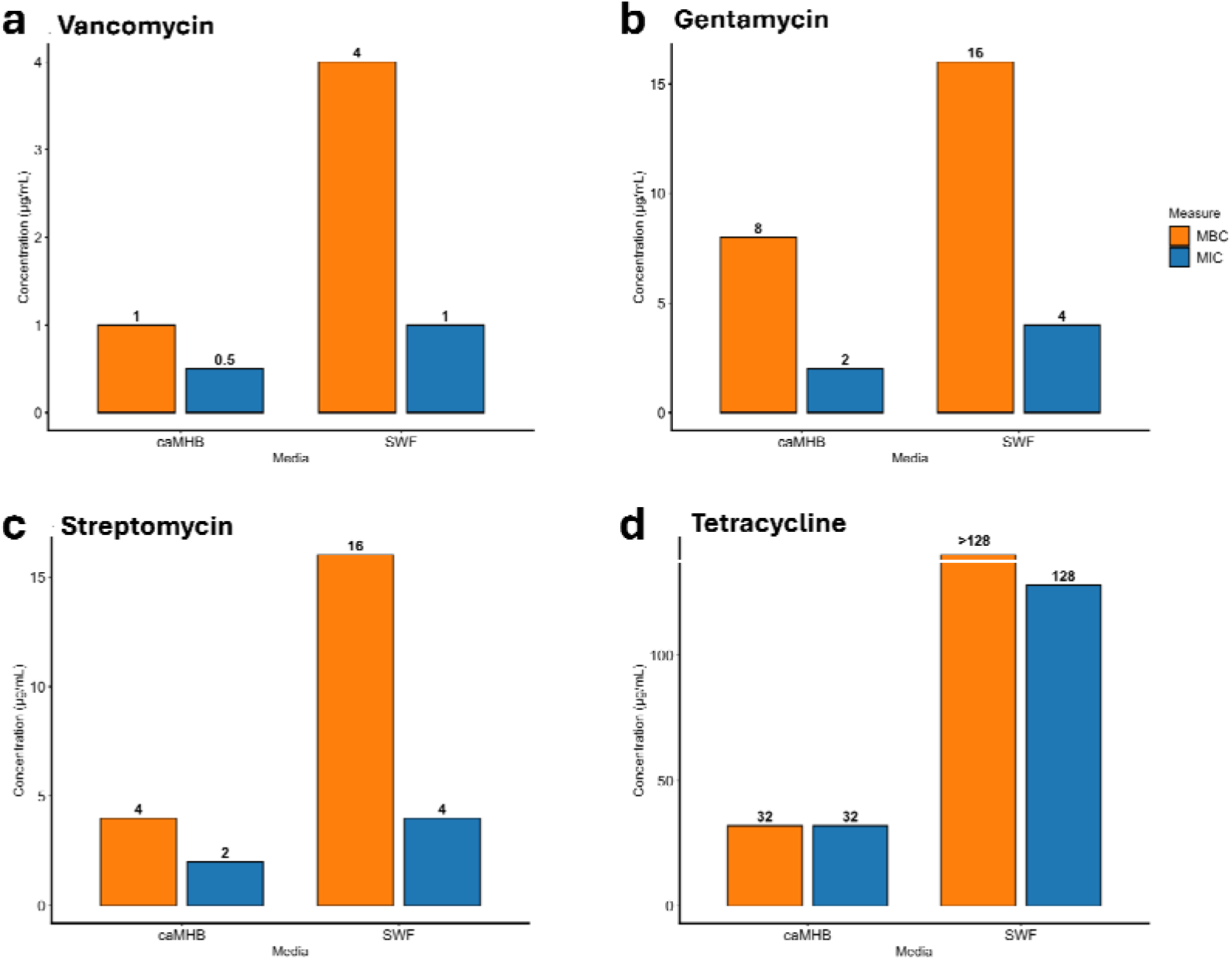
Higher antibiotic tolerance of MRSA in simulated wound fluid. Broth microdilution assay was used to assess susceptibility of *S. aureus* USA300 LAC to vancomycin **(a)**, gentamycin **(b)**, streptomycin **(c)**, and tetracycline **(d)** in both caMHB and SWF, and the MIC (minimum inhibitory concentration) and MBC (minimum bactericidal concentration) were recorded. Both MICs and MBCs were consistent across biological triplicates.

## Discussion

Many studies investigating bacterial pathogenesis and preclinical antimicrobial activity often make use of standard laboratory media, which do not provide an accurate representation of disease-specific complexities associated. This might be one of the contributing factors to the failure of many high-potential antimicrobials during clinical trials. This study revealed a strong variation in MRSA gene expression profile and antimicrobial susceptibility in two growth environments. We showed that the growth environment strongly influences the expression of genes associated with MRSA pathogenicity and physiological responses such as transmembrane transport.

In this study, we revealed that MRSA downregulates its expression of genes involved in transmembrane ion transport when growing in a wound-mimicking environment compared to the standard laboratory media, caMHB. Most notably among these genes are the potassium (*kdpFABC*) and phosphate (*pstSCAB* and *phoU*) transmembrane transporter genes. Potassium and phosphate are some of the major ions in the cell. The Kdp complex is a high-affinity ATP-dependent potassium transporter and has been previously reported to be highly overexpressed when *S. aureus* is grown in a media containing high sodium concentration (Price-Whelan et al., 2013). This potassium transport system helps to maintain a stable potassium gradient across the bacterial membrane by transporting potassium ions into the cell, as this is important in bacterial homeostasis (Zhang et al., 2023; Do and Gries, 2021). Hence, increased expression of this system in caMHB indicates a higher concentration of sodium ions in caMHB than in SWF and a possible disruption of the membrane gradient. The Kdp system has been implicated in bacterial pathogenesis. KdpDE has been previously reported to regulate virulence factors in *S. aureus,* including the capsular polysaccharide produced by many MRSA strains (Freeman et al., 2013; Zhao et al., 2010). The Pst phosphate transport system is a high-affinity phosphate transport system that helps the cell to transport phosphate from the periplasm across the plasma membrane (Jansson, 1998). This high affinity phosphate transport system is often activated during phosphate starvation. Perhaps, the significant upregulation of this pathway in caMHB (downregulation in SWF) indicates deficiency in intracellular phosphate in *S. aureus* USA300 LAC when grown in caMHB or a possible saturation of phosphate in SWF. In addition to its role in homeostasis, disruption (overexpression) of the PstSCAB-PhoU operon has been reported to attenuate S. aureus virulence in an in vivo mouse model by increasing S. aureus sensitivity to nutritional immunity (Kelliher et al., 2020). Additionally, during skin and soft tissue infections, antibacterial compounds such as nitric oxide are produce by the immune cells, the upregulation of PstSCAB-PhoU system helps *S. aureus* to survive under nitric oxide stress (Stephens et al., 2023). However, their role in *S. aureus* resistance to clinically-relevant antibiotics is not yet known. The variations in the expression of these transport systems in both media is an indication of differences in nutritional availability, which influences transmembrane transport activity and membrane potential stability. A stable membrane potential contributes to antimicrobial susceptibility and the differences observed in the membrane potential in these two media might as well be linked to the variations in antimicrobial susceptibility of MRSA in these two media.

Another interesting gene expression pattern seen in this study is the upregulation of two MFS multidrug resistant genes, *norB* and *norC*. NorB and NorC are known to help *S. aureus* pump out antibiotics and toxic compounds (Costa et al., 2013). The slight increase in antibiotic tolerance we observed in SWF might be linked to the increased expression of these efflux genes. Other studies have also reported variations in antibiotic tolerance in different growth media (Orababa et al., 2026; Bock et al., 2018). The role of these two efflux genes in wound environment, in the absence of antibiotics, is currently not clear, perhaps they are involved in pumping out some components of serum or associated with adapting the bacterial pathogen to this environment. It is also currently being hypothesised that some of these multidrug efflux pumps might be playing another role beyond just pumping out antibiotics and this may include virulence or biofilm formation (Whittle et al. 2024; Vareschi et al., 2025). Future studies should focus on understanding the roles of these multidrug efflux pumps in MRSA virulence and biofilm formation in chronic wound environment.

Our study also revealed the upregulation and downregulation of multiple genes in SWF compared to caMHB. One of the interesting groups of genes that were significantly upregulated are the genes coding for serine protease-like protease *splABCDEF*. These genes have been previously reported with *S. aureus* virulence and immune evasion (Pospich et al., 2026). Pospich et al., (2026) reported that SplB elicits T-cell activation and cytokine secretion in atopic dermatitis patients. This protease has also been associated with the inactivation of human compliment system by inactivating all three compliment pathways and blocking opsonophagocytosis (Dasari et al., 2022). As a result, monoclonal antibodies against SplB are now being considered for the development of new vaccines against *S. aureus* infections (Igbal et al., 2025). Fibronectin binding protein-coding genes (*fnbA* and *fnbB*) were also differentially expressed upregulated in SWF. The significant upregulation of these genes in wound-mimicking growth environment (SWF) aligns with their previously reported role in *S. aureus* skin colonisation and infection (Edwards et al., 2011; da Costa et al., 2022). These fibronectin-binding proteins are majorly used for cellular attachment and have been previously implicated in *S. aureus* skin colonisation and infection (Edwards et al., 2011; da Costa et al., 2022). For instance, FnbB has been previously shown to promote the binding of *S. aureus* to healthy corneocyte and loricirin, the primary component of the cornified cell envelops (da Costa et al., 2022). Perhaps their upregulation in SWF indicates their role is cell-to-cell attachment leading to the formation of micro-colonies/cellular aggregation as seen in MRSA biofilm in SWF and soft tissue wound model. This possibly accounts for the low biofilm biomass reported in the SWF using the crystal violet method. Due to the formation of cellular aggregates (in SWF) as opposed to attachment to the microtitre plate (as in the case with biofilm formed in caMHB), the washing step in the crystal violet method washed off many cellular aggregates, thus reducing the biofilm biomass in SWF. This further highlights some of the drawbacks in the use of general purpose media and models in accurately predicting *in vivo* bacteria activity.

There was also significant upregulation of genes coding for *S. aureus* toxins (*hla, hlgCB,* and *psm-*ZI) in SWF. There are evidence of these toxins mediating or promoting *S. aureus* pathogenesis in skin infections. ZI-hemolysin (Hla) is one of the most characterised *S. aureus* toxins and has been reported to mediate vascular injury in *S. aureus*-induced dermonecrosis (Yang et al., 2024). Both *S. aureus* Y-hemolysin (HlgCB) and psm-ZI are pore-forming exotoxins that have been linked with *S. aureus* pathogenesis (Pivard et al., 2023). The *psm*-ZI locus has also been reported to regulate the production of *hla*, with the deletion of either gene leading to similar virulence defect in *S. aureus* skin infection (Berube et al., 2014). The significant upregulation of these virulence genes indicates an increased virulence of *S. aureus* in the simulated wound environment compared to caMHB, supporting our hypothesis of the impact of environment on bacterial pathogenesis.

In conclusion, our work highlights the importance of the growth environment in bacterial physiological responses and susceptibility to antibiotics. Hence, it emphasises the need to use the right growth media and infection model in research that aims to better understand bacterial virulence. This is also important in preclinical testing of new antimicrobials as well as susceptibility testing during clinical diagnosis. This study has just provided an overview of the variations in gene expression between two growth environments; it has not fully provided the mechanistic explanation for some of these variations. Most especially the role of the Pst and Kdp ion transport systems in bacterial virulence and resistance to antibiotics. Future studies should focus on comparing the differences in gene expression between other commonly used standard growth media (such as the Luria Bertani broth and Tryptic Soy broth) in research and infection-mimicking media to understand how these environments might influence virulence and antibiotic susceptibility.

## Methods

### RNA sequencing and analysis

Overnight cultures of *S. aureus* USA300 were diluted to 0.1 OD_600_ in caMHB or SWF, grown for 6h, and diluted to 0.8 OD_600_. The cultures were washed by centrifuging at 10,000 rpm for 10 min. Cell pellets were then resuspended in PBS, centrifuged at 10,000 rpm for 10 min, snap frozen, and sent for RNA extraction and sequencing by Genewiz. Quality of the raw fastq files was checked with fastqc (Leggett *et al.,* 2013), after which trimommatic (Bolger *et al.,* 2014) was used to remove Illumina adapter sequences from the reads. Alignment was done with Bowtie2 (Langmead and Salzberg, 2012). All reads were aligned to *S. aureus* USA300 FPR3757 whole genome sequence (GCF_000013465.1). The aligned sam files were converted to bam format using samtools (Li *et al.,* 2009), after which featureCounts (Liao *et al.,* 2014) was used to obtain read counts. DESeq2 (Love *et al.,* 2014) in R v4.3.3 (R Core Team, 2023) was then used to determine differentially expressed genes, while Gene Ontology (GO) analysis was performed with ShinyGO (Ge et al., 2019).

### Membrane disruption assay with DiSC3(5)

A DiSC3(5) (3,3’-Dipropylthiadicarbocyanine iodide) assay was performed as previously described by Buttress et al. (2022) with a little modification. Briefly, *S. aureus* USA300 LAC bacterial culture growing at the exponential phase in LB was diluted to 0.5 OD_600_ with PBS containing 0.5 mg/ml bovine serum albumin (BSA) (Sigma-Aldrich, UK) and 2 µM glucose. The addition of BSA was to prevent the binding of DiSC3(5) reagent to the microtitre plate. Cells were washed in PBS containing 0.5 mg/ml bovine serum albumin (BSA) (Sigma-Aldrich, UK) and 2 µM glucose twice and then resuspended. After resuspension, the tube containing washed cells was incubated for 15 min. Bacteria were then transferred into a black polystyrene 96-well plate (Corning Inc., US) and autofluorescence measured for 5 min in Tecan SPARK 10 M (Tecan, Switzerland) plate reader, after which an equal volume of 1 µM DiSC3(5) reagent (Invitrogen) in 1% DMSO was added to give a final DiSC3(5) concentration of 0.5 µM and fluorescence measured (excitation wavelength = 610±10, emission wavelength = 660±10) for another after 1h in a Tecan SPARK 10 M (Tecan, Switzerland) plate reader.

### Antimicrobial Susceptibility testing

The broth microdilution method was used to determine the minimum inhibitory concentration of the antimicrobials as recommended by the European Committee on Antimicrobial Susceptibility Testing (EUCAST). Briefly, the bacterial strain of interest was streaked on LB agar and incubated at 37°C for 18-24 h to produce distinct colonies. Twice the maximum concentration of the antibiotics to be used was prepared in the medium of interest (caMHB, SWF) and dispensed (100 µl) in the first column of a Corning Costar CLS9018 (Corning Inc., US) 96-well plate. Fifty microlitres of the medium were then dispensed into each of the other wells, after which a two-fold serial dilution was carried out. A bacterial suspension in the medium was prepared by touching 3-4 distinct colonies with a sterile cotton swab and dispersing them in PBS. This was then standardised to a 0.5 MacFarland standard with a spectrophotometer (OD600nm = 0.08-0.10). The standardised bacterial suspension (50 µl) was then inoculated into triplicate wells of each concentration except for the sterile control well (without antimicrobials and bacteria). Growth control wells (with only bacteria) were also set up. The 96-well plates were sealed with Parafilm^TM^ and incubated at 37°C for 18-24 h. The lowest concentration well with no observable growth is taken as the MIC. After incubation, an inoculum of 10 µl was spotted out onto LB agar from each of the wells to determine the minimum bactericidal concentration (MBC). The LB agar plates were incubated at 37°C for 18-24 h. The concentration with no growth on the LB agar was taken as the MBC. Uninoculated media (without antibiotics) were used as a negative control, while bacterial suspension without antibiotics was used as a positive control.

### Biofilm assay

Biofilm biomass was assayed using crystal violet staining. Briefly, USA300 was streaked on LB agar and incubated at 37°C for 18-24 h to produce distinct colonies. Colonies were suspended in CaMHB and SWF. They were both adjusted to 0.5 MacFarland standard (OD600nm = 0.08-0.10) after blanking using fresh uninoculated medium. A 24-well (Corning Costar, USA) plate was prepared by filling three wells per media type with 2 mL of bacterial suspension. The remaining wells were filled with sterile media as contamination controls. The plate was incubated at 37°C without shaking for 48 hours. After incubation, planktonic cells and floating aggregates were transferred to a new plate. The remaining attached biofilms were washed twice with PBS to remove unbound cells. The attached biofilm was stained with 0.01% crystal violet for 15 minutes, then washed 3 times with sterile water. The plate was dried for 30 minutes under the laminar flow to remove excess water. Thereafter, the bound crystal violet was solubilized using 30% acetic acid, and the biofilm biomass was quantified by measuring absorbance at 590 nm.

### Calcofluor white staining of soft tissue wound biofilm

The *in vitro* soft tissue wound was made as previously described by Werthén et al. (2012) with some modification. In summary, bacterial suspension of *S. aureus* USA300 in exponential phase of growth was diluted to 0.08-0.1 OD_600_ (approximately 10^8^ CFU) and used to infect a thin film of collagen wound matrix on sterile glass slide (prepared as described above). This was incubated at 37°C for 24 hours to allow biofilm formation. After 24 h, calcofluor white (100 µl of original concentration) (Sigma-Aldrich, USA) was added to the slides and incubated for 15 minutes at 37°C. The slides were then fixed with 4% formaldehyde, washed three times with PBS and viewed using a confocal laser scanning microscope (Zeiss LSM 880) with the oil immersion magnification (x100) lens.

## AUTHOR CONTRIBUTIONS

Author contributions following CRedit Taxonomy:

Conceptualisation: OQO

Investigation: OQO, AK, LIU

Formal analysis: OQO, AK, LIU

Methodology : OQO, AK, LIU

Supervision: OQO

Writing – original draft: OQO

Writing – review & editing: all authors.

## ACKNOWLEDGMENTS

This work benefited greatly from feedback and discussion with Freya Harrison. We also gratefully acknowledge the support of the University of Warwick Institute of Advanced Studies (IAS) through an Early Career Fellowship awarded to OQO during the writing up of this manuscript.

## ETHICS APPROVAL

No ethical approval was required for this study.

